# Decoding voluntary movements and postural tremor based on thalamic LFPs for closed-loop stimulation for essential tremor

**DOI:** 10.1101/436709

**Authors:** Huiling Tan, Jean Debarros, Alek Pogosyan, Tipu Z. Aziz, Yongzhi Huang, Shouyan Wang, Lars Timmermann, Veerle Visser-Vandewalle, David J. Pedrosa, Alexander L. Green, Peter Brown

## Abstract

High frequency deep brain stimulation (DBS) targeting motor thalamus is an effective therapy for essential tremor (ET). However, conventional continuous stimulation may deliver unnecessary current to the brain since tremor mainly affects voluntary movements and sustained postures in ET. We recorded LFPs from the motor thalamus, surface electromyographic (EMG) signals and/or behavioural measurements in seven ET patients during temporary lead externalization after the first surgery for DBS when they performed different voluntary upper limb movements and in nine more patients during the surgery, when they were asked to lift their arms to trigger postural tremor. We show that both voluntary movements and postural tremor can be decoded based on features extracted from thalamic LFPs using a machine learning based binary classifier. This information can be used to close the loop for DBS so that stimulation could be delivered on demand, without the need for peripheral sensors or additional invasive electrodes.

## Introduction

Continuous high frequency deep brain stimulation (DBS) targeting ventral-intermediate thalamus is an effective therapy for medically refractory essential tremor (ET) [1–4]. However, as many as 70% of patients lose the benefit of DBS over time [5], due to disease progression or habituation to stimulation [6]. These circumstances often require an increase in the energy delivered through increased amplitude, frequency, and/or increased pulse width, which is commonly associated with more pronounced adverse effects of stimulation, resulting in slurred speech, unpleasant sensations, incoordination and walking difficulty [7]. Furthermore, tremor in ET is typically intermittent, predominantly occurring during voluntary movements and sustained postures [8–10], suggesting that stimulation could be more focussed in time to control symptoms.

Closed-loop stimulation, in which stimulation parameters are automatically adjusted to stimulate on demand, is seen as a potential breakthrough for the treatment of essential tremor [2,11]. Prior studies of closed-loop DBS for ET have used wearable inertial sensors [12,13] and/or surface electromyography (EMG) [14] to provide feedback for the control of the stimulator. While wearable sensors provide reliable measurements of tremor, the wireless communication between the pacemaker and the external sensors introduces a potential vulnerability to the system due to breaks in transmission or hacking. In more recent studies [15,16], movement-related signals have been recorded using an extra strip of electrodes chronically implanted over the surface of the motor cortex. These signals are used to detect movements and to activate the DBS [8]. A recent single case study has shown that this approach can reduce tremor during writing and spiral drawing [16]. Nevertheless, the use of cortical strip electrodes introduces further instrumentation, risk of increased surgical complications such as haemorrhage, and additional cost [17]. In addition, detection of movement to trigger DBS may not provide sufficient control of tremor during sustained posture which is also an important aspect of ET [9].

Local field potentials (LFPs) recorded in the motor thalamus contain information related to both voluntary movement and postural tremor. Movement-related potentials in ventral intermediate (ViM) thalamic LFPs were observed with a similar latency as in the cortex [18,19]. In the frequency domain, reduction in the power of beta oscillations (14-30 Hz) and increase in a broad gamma frequency range (55-80 Hz) were reported in ViM thalamus during movements [18,20]. Ventral thalamic nucleus also expresses activities relating to ongoing tremor. For example, populations of neurons in the ViM thalamus exhibit tremor-frequency activity during tremor but not during rest [21]. Increased synchronisation at tremor and double tremor frequency in the ventral lateral posterior (VLP) nucleus of the thalamus has been associated with the presence of tremor [22–24].

Here we show that both voluntary movements and postural tremor can be decoded based on LFPs recorded from the same electrodes that are implanted in motor thalamus for therapeutic DBS. Importantly for the practical application of the proposed methods in decoding movements and triggering DBS, the classifier identified during training sessions based on data recorded while patients performed pre-defined movements can also be used to decode different natural movements such as drawing and pointing. Additionally, we show that postural tremor can also be decoded but the features for decoding postural tremor are different from those for decoding movements, suggesting a separate model would be required in order to deliver stimulation during postures that provoke tremor without further voluntary movements. Together these results pave the way for adaptive DBS based on LFP signals recorded from motor thalamus, without additional invasive electrodes or external sensors.

## Results

### Activities in ViM thalamic LFPs are modulated by movements

LFPs were recorded from seven ET patients when they performed different voluntary upper limb movements (Table 1 and Fig. 1). Average time-evolving power spectra of changes in ViM thalamic LFPs induced by movements were derived by aligning the normalized power spectra to all contralateral movement onsets and averaging across all individual movements in an experimental run. In this way, several frequency bands were identified over which increases or decreases in power could be distinguished during movements when averaged across trials (Fig. 2). These were a power increase in the theta/alpha band (4-12 Hz), power reduction in the beta range (13-34 Hz), and power increases in the mid gamma (56-95Hz) and high-gamma/high-frequency (105-195 Hz) ranges. However, the peak frequencies and ranges of movement-related changes varied from patient to patient.

**Figure 1.**
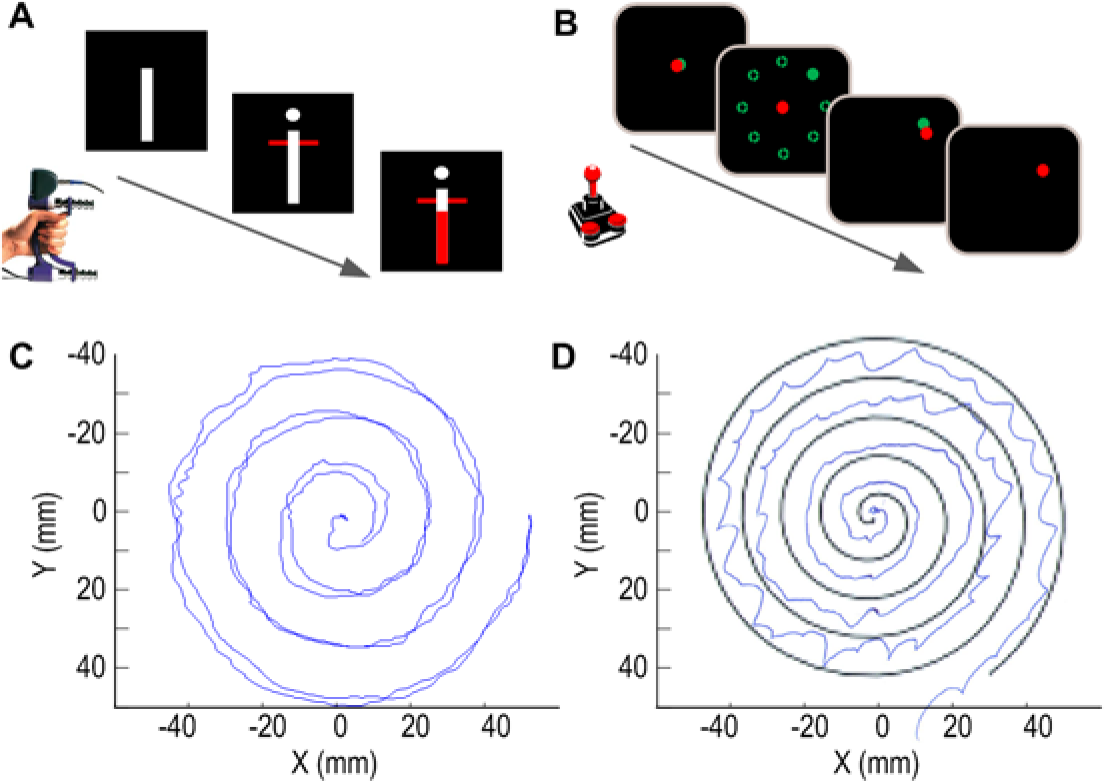
Examples of voluntary movements tested in the study. A) Cued hand gripping movements in 3 patients; B) Finger joystick movements in 1 patient with both hands separately; C) and D) Self-paced spiral drawing used for cross-task validation in 4 patients. Spiral drawing also revealed variable degrees of tremor in the immediate post-operative period.

**Figure 2.**
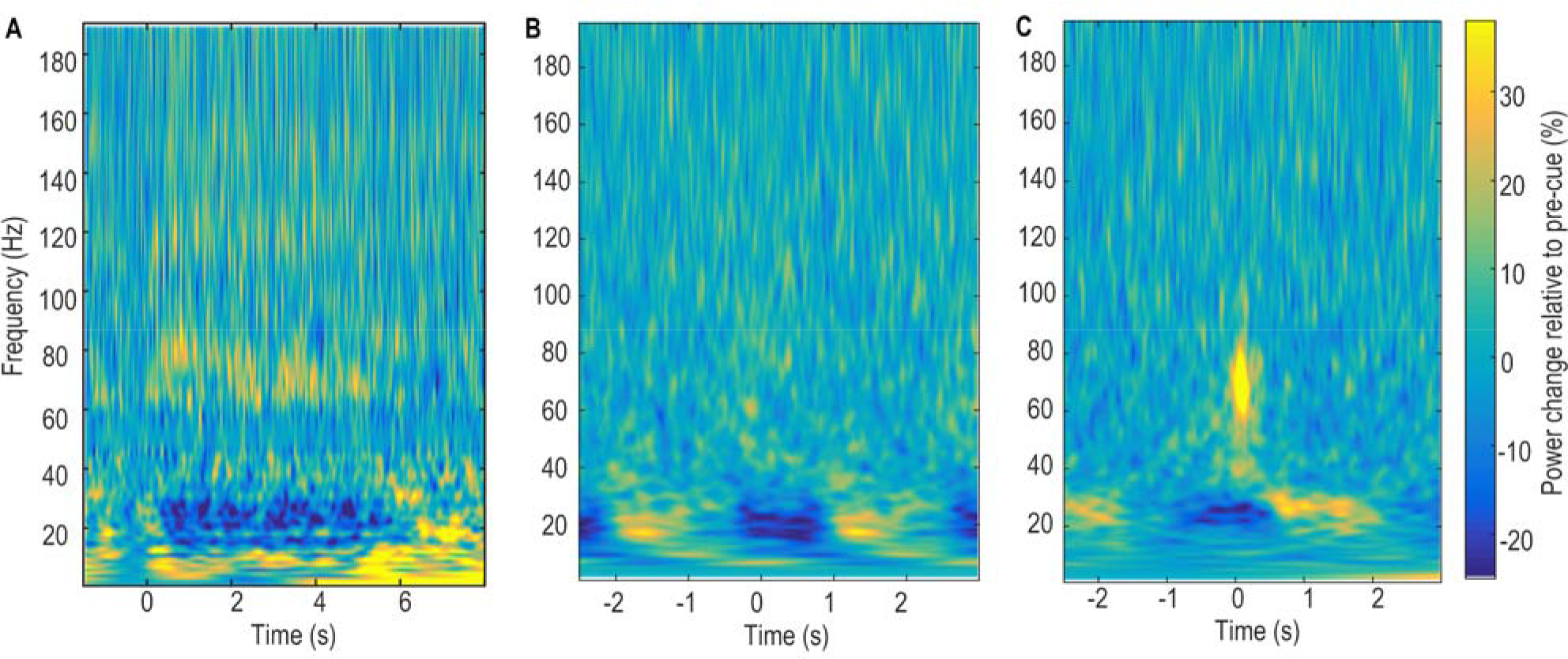
Examples of power changes induced by movements in ViM thalamus contralateral to the voluntarily moved hand. Changes were quantified relative to the average of the whole recording session and averaged across all trials during A) hand gripping for patient Ox1; B) finger joystick movements for patient Ox2; C) button pressing for patient Ox3. Time zero represents onset of individual movements.

**Table 1.** Patient details and motor tasks that have been tested. Patients in Oxford (Ox**) were recorded post-operatively during temporary lead externalisation and different motor tasks have been tested. Patients in Cologne (Cl**) were recorded inside the theatre during the surgery.

| ID | Age | Gender | Recorded hemisphere | Recording electrode | Pre-defined movements | Cross Task Validation |
| --- | --- | --- | --- | --- | --- | --- |
| Ox 1 | 35 | M | Left ViM | 3389; Medtronic | Cued Gripping Force (right hand only) |  |
| Ox 2 | 54 | F | Bilateral ViM | 3387; Medtronic | Cued Joystick Movement (both hands separately ) |  |
| Ox 3 | 62 | F | Left ViM | 3387; Medtronic | Cued button pressing (right hand only) | Drawing (right hand only) |
| Ox 4 | 26 | M | Bilateral ViM | 3387; Medtronic | Self-paced continuous finger tapping (both hands) | Drawing (both hands separately) |
| Ox 5 | 37 | F | Left ViM | 3387; Medtronic | Self-paced continuous flexion /extension of right wrist | Drawing (right hand only) |
| Ox 6 | 65 | F | Bilateral ViM | 6180; St. Jude Directional | Cued Gripping Force (both hands separately) | Self-paced reach and grasp + Self-paced pegboard movement (right hand only) |
| Ox 7 | 74 | M | Bilateral ViM | 3389; Medtronic | Cued Gripping Force + Self- paced continuous tapping (both hands separately) | Drawing + Self-paced pegboard movement (right hand only) |
| Cl 1 | 64 | M | Right VLp | micro-macroelectrode (LFPs recorded from the macro contacts were used for analysis) | Keep arm rest for 30-60 s and then elevate and hold their forearm at an angle of ~30° and to spread their fingers for 30–60 s |  |
| Cl 2 | 52 | M | Right VLp |  |  |  |
| Cl 3 | 75 | F | Left VLp |  |  |  |
| Cl 4 | 71 | F | Left VLp |  |  |  |
| Cl 5 | 73 | F | Bilateral VLp |  |  |  |
| Cl 6 | 67 | F | Bilateral VLp |  |  |  |
| Cl 7 | 69 | F | Bilateral VLp |  |  |  |
| Cl 8 | 62 | M | Left VLp |  |  |  |
| Cl 9 | 72 | M | Bilateral VLp |  |  |  |

### Both cued brief movements and blocks of continuous movements are detected

The activities over 8 different frequency bands in the LFPs recorded from the contralateral ViM thalamus was quantified: 1-3 Hz, 4-7 Hz, 8-12 Hz, 13-22 Hz, 23-34 Hz, 35-45 Hz, 56-95 Hz and 105-195 Hz. These features and the history of these features over a 1 second window with 100 ms update interval were used to decode whether the patient was engaged in voluntary movement at each time point using a logistic regression (LR) based binary classifier. The within-task cross-validation tests showed that ViM LFPs could be used to detect hand gripping despite the variation in force generated in each grip (Fig. 3 - Figure Supplement 1A), as well as joystick movements (Fig. 3 - Figure Supplement 1B) or button pressing (Fig. 3 - Figure Supplement 1C) on individual trial basis, despite the short duration of individual movements. The same approach could also detect blocks of self-paced continuous movements (Fig. 3 - Figure Supplement 2). The LR-based classifier output consistently increased when movements occurred and remained high until the movement stopped.

The ROC curves (Fig. 3A) show that decoding of movements was achieved well above chance-level in all the patients and for the different types of movements recorded. The AUC ranged between 0.74 and 0.89 for cued brief movements, and between 0.89 and 0.99 for blocks of continuous movements. With a constant threshold of 0.4, 95.6% ± 2% (mean ± SEM across different test session) of individual movements were detected with a mean latency of −300 ms. The negative detection latency meant that the LR-based classifier output exceeded the decision threshold 300 ms before the actual movement onset. With the decision threshold of 0.4, the decoding sensitivity was between 0.67 and 0.84 for brief movements and between 0.76 and 0.99 for continuous blocks of movements. The corresponding false positive rate was between 0.15 and 0.33 for brief movements, and between 0.002 and 0.20 for continuous movements. This meant that if movement detection was used to actuate DBS, the DBS would be switched on 80.8% ± 2.6% of the time when the patients were making any voluntary movements, and the DBS would be switched on 20.0% ± 3.0% of the time when the patients were at rest.

**Figure 3.**
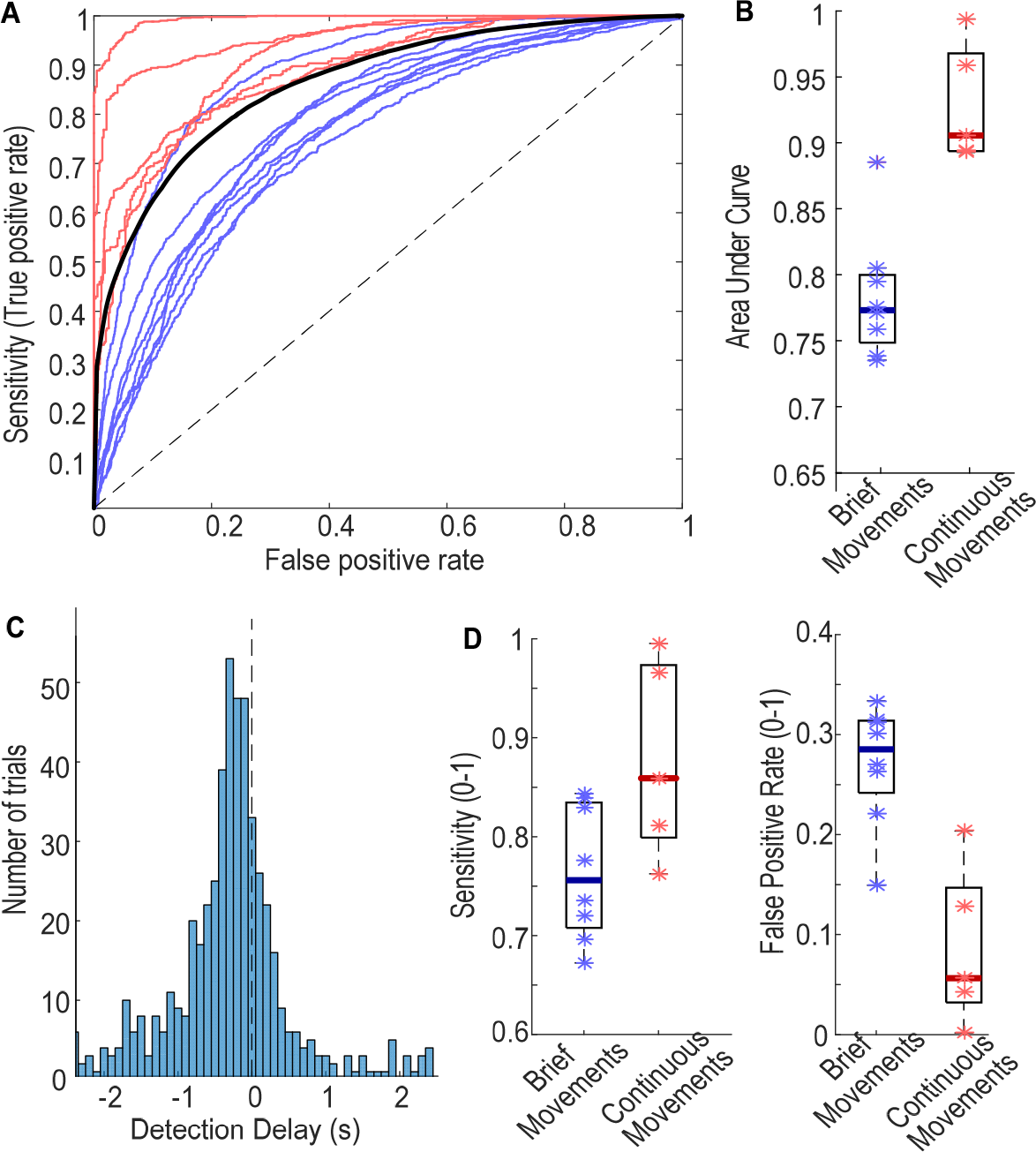
Evaluation of movement detection. A) ROC plots showing sensitivity against the false positive rate for the different possible thresholds used for decoding. Blue and red thin lines show the ROC curves of individual cases for cued brief movements and self-paced continuous movements, respectively. The thick blue line shows the ROC averaged across all cases. B) Area Under the Curve (AUC). C) Histogram of detection delays of individual movements. Zero is movement onset. D) Sensitivity and false positive rate for detecting brief and continuous movements, respectively, with a constant decision threshold of 0.4.

### Contribution of different ViM LFP features in decoding of movement

Fig. 4A shows the contribution of LFP features in each frequency band and at each time lag for movement decoding averaged across all test recording sessions. Activities in the low beta band (13-22 Hz) contributed most to the movement decoding, and this was followed by activities in the theta (4-7 Hz), delta (1-3 Hz), alpha (8-12 Hz), and high beta band (23-34 Hz) in order of contribution (Fig. 4B). The decoding performance remained high after removing activities lower than 8 Hz, which could potentially be contaminated by movement artefacts, from the Logistic regression. Similar decoding performance was reached after further removing activities higher than 45 Hz, which might be contaminated by stimulation artefact if DBS were switched on. However, if only the broad-band beta activity and its history were included, the decoding performance was noticeably lower (Fig. 4C).

**Figure 4.**
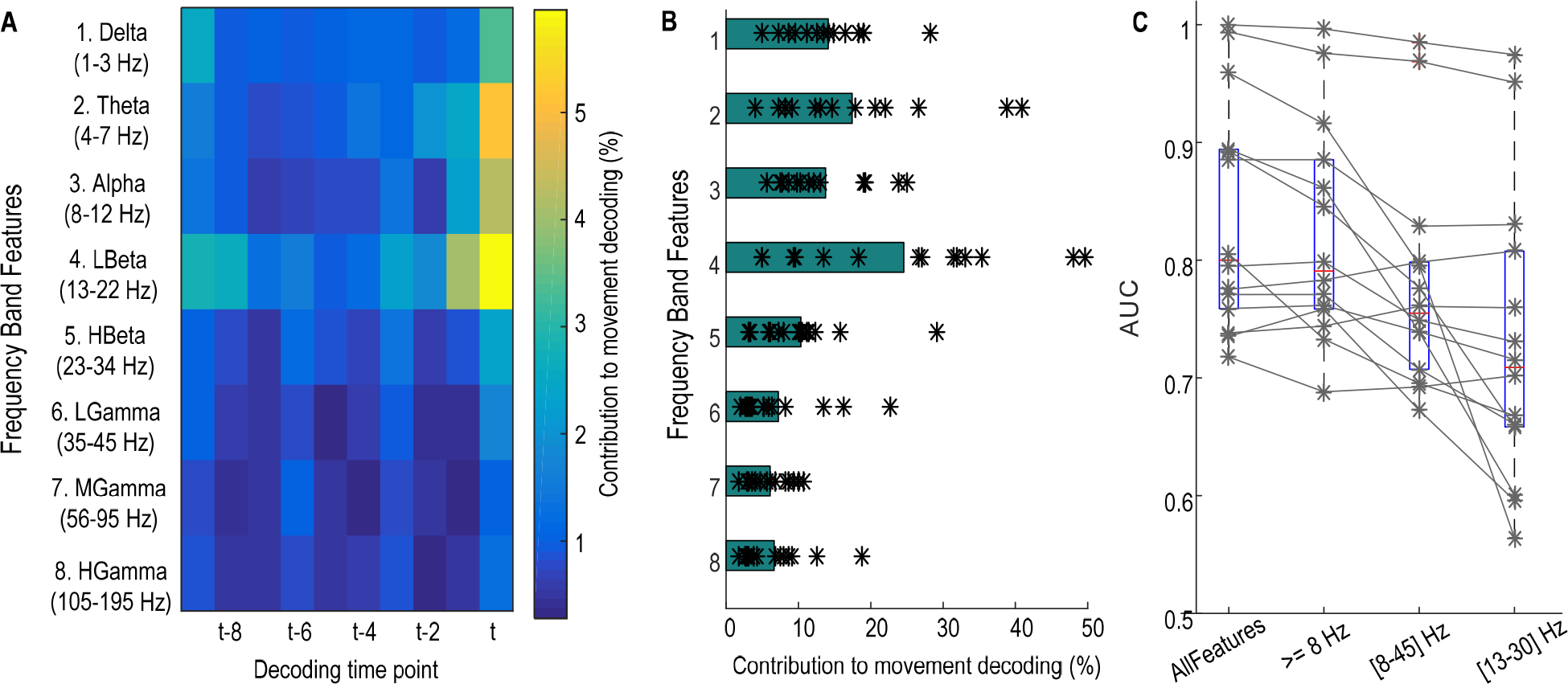
Features important for movement decoding. A) Weight (absolute value) distribution across features indicating low beta activities (13-22 Hz) at (t) and just before (t-1) movement onset contributed most to decoding movements; B) Sum of weights for features of different frequency bands, ordered as in A. C) AUC values were not much affected by removing features lower than 8 Hz. AUC values when considering only 8-45 Hz, and only beta band activities (13-34 Hz) are also shown.

### Cross-task validation of movement detection

The LR-based classifier trained using data recorded while the subjects performed pre-defined cued movements was used to decode other self-paced voluntary movements such as drawing, reaching and picking up objects (Fig. 5). In all the 8 cross-task validation test sessions from 5 patients, the AUC of the movement detection was 0.82 ± 0.023. With a constant threshold of 0.4, the sensitivity for movement detection was 0.77 ± 0.038 and the false positive rate was 0.23 ± 0.033. This meant that if movement detection were used to actuate DBS, DBS would be switched on 77% ± 3.8% of the time when the patients were engaged in free voluntary movements, and DBS would be switched on 23% ± 3.3% of the time when the patients were at rest.

**Figure 5.**
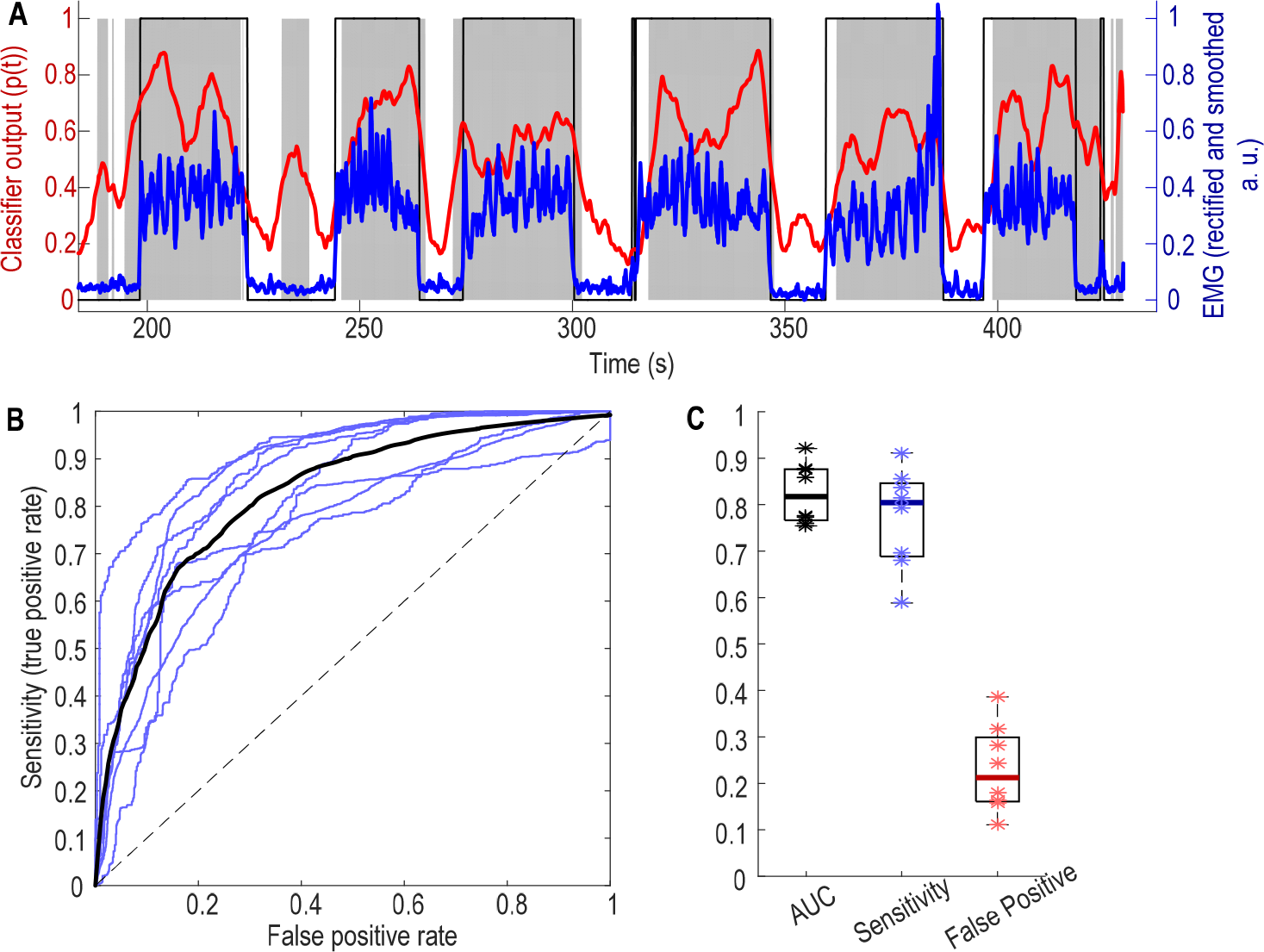
Results of cross task validation test. A) Models trained based on data recorded when the patient performed cued hand gripping can detect self-paced pegboard movement in the same patient (Ox6). Blue lines show normalized EMG measurements; thin black lines show time points with movement labelled according to EMG; thick red line shows output from the classifier, and shaded grey areas show the time points with decoded movement probability larger than 0.4. B) ROC of eight individual cross-task test sessions in blue from 5 patients and average across all test sessions in black; C) AUC values, sensitivity and false positive detection rate at the threshold of 0.4. * indicates values from individual test sessions, thick horizontal lines show the average across all test sessions.

### Postural tremor detection

Tremor was considerably improved in the patients recorded in Oxford in the post-operative period, but one of the patients recorded post-operatively (Ox7) still displayed significant postural tremor when he was holding the arms abducted, up in the air with elbows flexed and the fingers of both hands pointing towards each other. Tremor disappeared when the patient stretched out both arms with both elbows straight. The postural tremor was evident in the increased 3-7Hz activity from the accelerometer attached to the hand. This postural tremor could also be decoded based on ViM LFP measurements, as shown in the results from the within-session cross-validation approach (Fig. 6A). The AUC for detecting tremor in different postures was 0.88. With the decision threshold of 0.4, the sensitivity of the detection was 80% and false positive detection was 22%. For the seven blocks of postural tremor recorded, the detection on average anticipated tremor onset by −0.1± 0.13 second, ranging from − 0.4 to 0.3 second. However, the LR model for detecting postural tremor, as represented by the weights attributed to different features (Fig. 6C), was very different from that optimised for decoding voluntary movements (Fig. 6D). Accordingly, the classifier optimised for decoding movement failed to decode postural tremor and had an AUC of 0.48, close to random decoding, indicating that separate models might be required to detect voluntary movements and postural tremor in the same subject.

**Figure 6.**
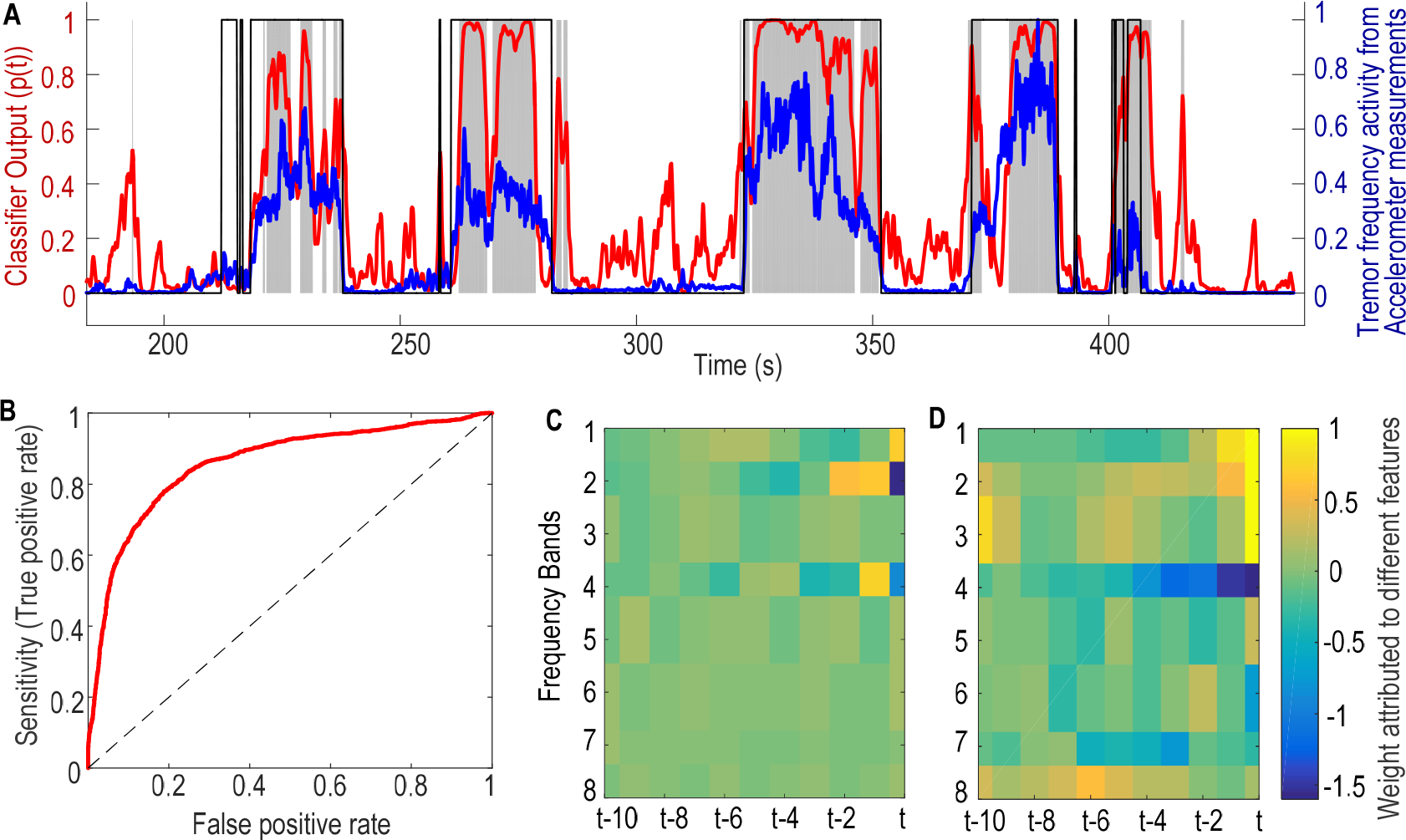
Postural tremor detection in a representative case (Ox7) A) Tremor displayed in different postures can also be detected based on ViM LFPs. The blue line shows the normalised tremor frequency power in accelerometer measurements; thin black line shows the time points with tremor judged from accelerometer measurements; the red line shows the classifier output based on ViM LFPs; grey shaded area show the time points with classifier output larger than 0.4. B) ROC plot of the tremor detection shows that 80% detection rate can be achieved with 20% false positive rate. The model optimised for tremor detection (shown in C) is very different from the model optimised for movement detection (shown in D) in the same subject. Feature labelling and time resolution is the same as in Figure 3.

In all the patients recorded intraoperatively in Cologne, postural tremor emerged after the elevation of the arm as shown by increased 4-7Hz activity in the EMG. The EMG activity during postural tremor was clearly differentiable from that at rest without tremor, with increased activity in the tremor and double tremor frequency bands (Fig. 7A). Based on the power of the tremor frequency activity in the EMG measurements, tremor was detected for 93% ± 2.9% of the time when the hand was elevated; in contrast, tremor was detected only for 5.4% ± 1.7% of the time when the hand was at rest. Postural tremor was also associated with increased activity in the tremor frequency band (4-7 Hz) in the thalamic LFPs (Fig. 7B). The LR-based classifier using features extracted from thalamic LFP as model inputs was used for detecting postural tremor in the contralateral limb. The ROC curves (Fig. 7C) showed that the decoding of postural tremor was achieved well above chance-level in all the 12 tested hands from the 9 patients. The AUC of tremor detection was 0.79 ± 0.027 (ranging between 0.66 and 0.96). With a constant threshold of 0.4, the sensitivity for movement detection was 0.77 ± 0.020 (ranging between 0.71 and 0.95) and the false positive rate was 0.29 ± 0.038 (ranging between 0.086 and 0.49). The oscillatory activities between 4-7 Hz (theta frequency band) in thalamic LFPs contributed most to the tremor decoding, and the AUC of the decoding increased with increasing levels of theta band modulation in thalamic LFPs relative to rest across tested hands (Spearman correlation, r12=0.825, p = 0.0017).

**Figure 7.**
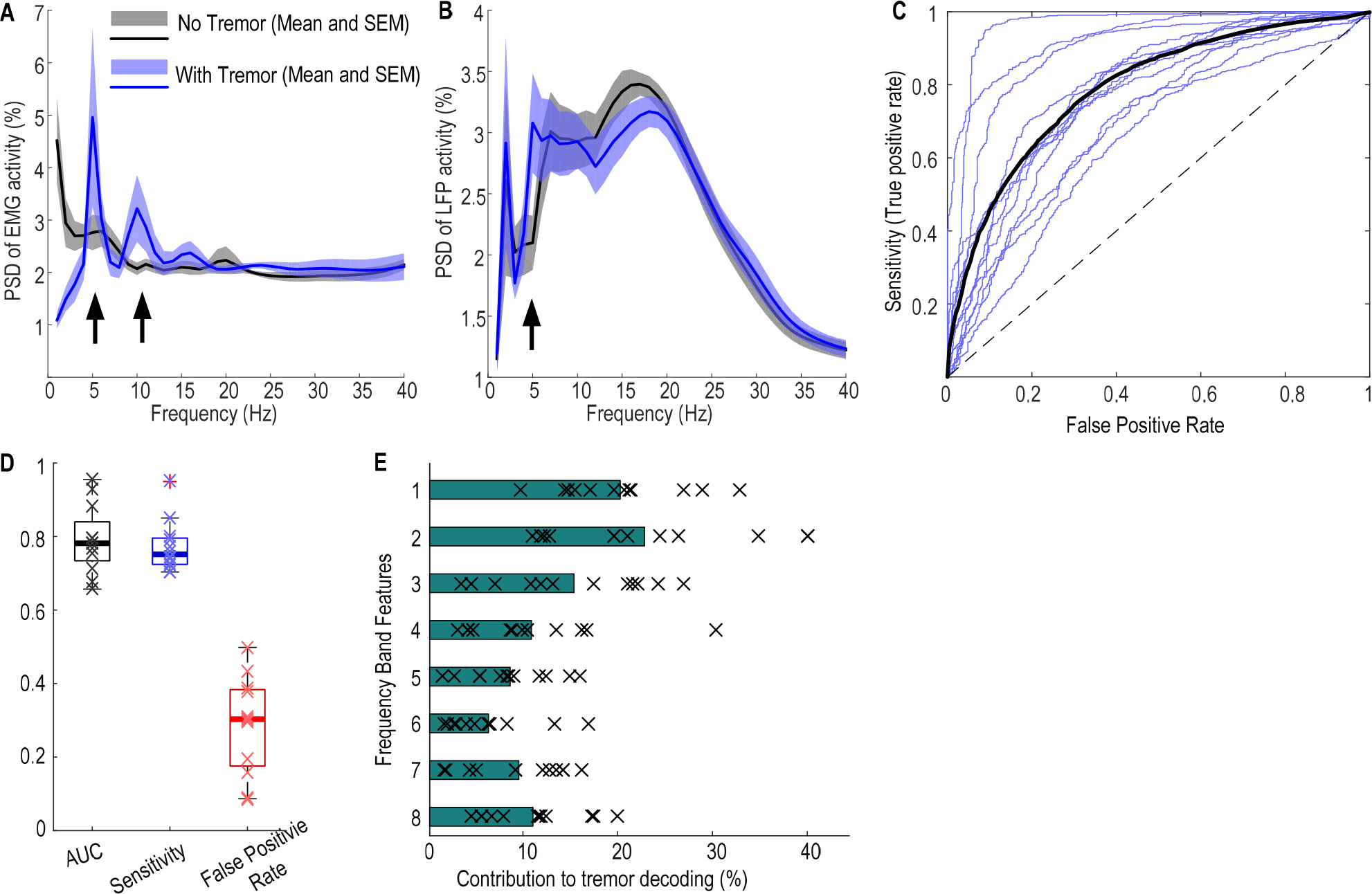
Postural tremor detection across subjects. A. Postural tremor was differentiated from rest by increased tremor frequency and double tremor frequency activities (indicated by the arrows) in the EMG; B. Postural tremor was associated with increased tremor frequency activities in the thalamic LFPs (indicated by the arrows). C. ROC curves of tremor detection based on thalamic LFPs (individual sides in thin blue lines and average across all sides in black). D. AUC, Sensitivity and false positive rates. Thick horizontal lines show the mean values and x show data from each individual test session. E. Activities between 4-7 Hz contributed most to the tremor decoding, with feature labelling the same as in Figure 3.

## Discussion

We have demonstrated that both voluntary movements and postural tremor can be detected based on thalamic LFPs recorded using the same electrode as used for therapeutic stimulation, with an average sensitivity of 0.8 and false positive rate of 0.2. Oscillatory activities in the low beta (13-22 Hz) and theta (4-7 Hz) frequency bands contributed most to detecting voluntary movements and postural tremor, respectively. The movement detection on average preceded movement onset. Critically, the same classifier trained on data recorded during prompted pre-defined movements was also able to detect different self-paced movements, representative of those made during everyday life. However, separate models are required for detecting voluntary movements and postural tremor to ensure that both situations will be detected to trigger the DBS.

### Implications for Closed Loop DBS for Essential Tremor

This study suggest that thalamic LFPs can be sufficient to trigger anticipatory DBS to suppress tremor during action and sustained posture. A previous study monitoring natural hand movements made during everyday life in healthy subjects showed that the hand was essentially at rest for approximately half the time when subjects were awake [25]. Accordingly, actuating DBS only during movement or during postural tremor could lead to up to 50% reduction in the total energy delivered to the brain during awake hours and possibly more once sleep is considered. Compared to previous studies [12–16], our results showed that responsive DBS for essential tremor can be achieved without the requirement of external sensors or additional electrocorticography strips. Using LFP activities recorded from the stimulation electrode for closing the loop for DBS has advantages in minimising the time delays and data loss associated with wireless communication with limb-mounted external sensors, and in minimising the surgical risk of additional invasive instrumentation.

Patients with ET may also develop tremor during sustained postures such as holding an open book. Tremor under these circumstances might not be addressed by only triggering DBS by voluntary movements, whether using thalamic LFPs or electrocorticographic recordings. In addition, decoding failed to detect voluntary movement in a small fraction of active movement trials. So an important aspect of the present study is the ‘failsafe’ procedure of detecting tremor should it develop. In contrast, and to our knowledge, there is still no evidence showing that postural tremor can be detected from cortical signals alone.

Nevertheless, there are a few important technical considerations related to using thalamic LFPs for closed-loop DBS. First, all results presented here are based on recordings made with stimulation switched off. Stimulation artefacts may lower the signal to noise ratio of LFPs recorded when stimulation is on. However, at least the detection of movement or tremor onset to start stimulation will not be affected by stimulation artefact. Yet once stimulation is switched on, the classifier still needs to detect the offset of movement or tremor to switch off stimulation, and here the performance of the classifier at this point in time may be compromised by the presence of stimulation artefact. Noteworthy, activities in the beta and theta frequency bands recorded from the stimulation electrodes contributed most to movement and tremor detection, respectively, and can both be monitored even during stimulation, with sufficient filtering and signal processing [26–29]. It remains to be seen whether a separate model with different weight vectors may be required for detecting movement and tremor offset based on LFPs recorded with simultaneous simulation. Second, in the approach proposed here, the sensitivity (here defined as the percentage of time when the stimulation would be on during the total duration of volitional movements) and false-positive rate (here defined as the percentage of time when the stimulation would be on during total time when this is not necessary) are dependent on the detection threshold. It is important to consider what is the desired sensitivity and false-positive rate for the best patient outcome in clinical practice. The detection rates and sensitivity results reported here were based on a constant threshold of 0.4 for all patients. This threshold could be further optimised for each patient according to factors such as tolerance to side effects and desirable levels of power saving. Third, in the patient who demonstrated significant remaining postural tremor in the post-operative period and for whom we also recorded voluntary movements, we showed that the model optimised for detecting postural tremor was very different from that optimised for detecting voluntary movements. Separate models for detecting movement and for detecting postural tremor may be required to ensure that DBS is actuated when intention and/or postural tremor is detected for optimal treatment of the disease. Considering all these issues, we propose the framework shown in Fig. 8 to detect both movements and tremor based on ViM LFPs for closed-loop DBS for ET.

**Figure 8.**
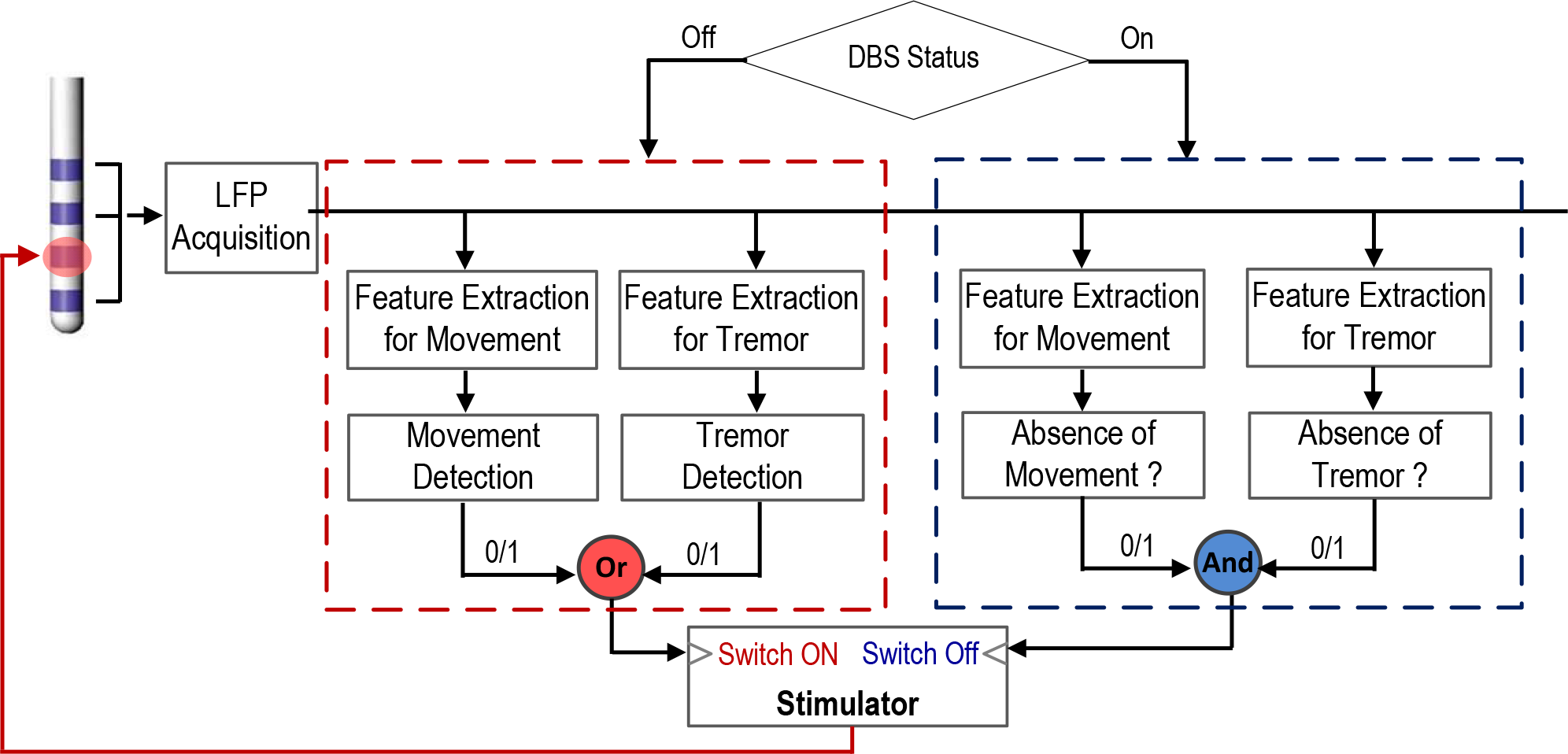
A proposed schematic for closed-loop DBS for Essential Tremor based on ViM LFPs. The detection of movement or tremor is used to actuate the DBS; and the detection of movement offset is used to switch off the DBS, but only provided tremor detection, if present, has ceased.

### Machine learning based approach vs. Threshold-based decision-making based on a single feature

In this study, we adopted a machine learning approach using a linear combination of activities in eight non-overlapping frequency bands in the thalamic LFPs for decoding. Even though we demonstrated that oscillatory activities in the low beta (13-20 Hz) and theta (4-7 Hz) frequency bands contributed most to the decoding of voluntary movements and postural tremor, respectively, activities in other frequency bands also contributed and increased the decoding accuracy. Thus the present approach may potentially be more effective than threshold-based closed-loop DBS based on activities in a single frequency band, as used in most previous studies [15,16,26]. This is because machine learning based algorithms can automatically attribute appropriate weights to multiple features specific to each patient through optimisation. The machine learning algorithm used in this study was based on a linear regression model which can be trained with data recorded over just a few minutes, and can also be easily implemented in real-time for closed-loop DBS applications.

### Limitations and Caveats

A few general caveats should be borne in mind. First, in the data analysed for decoding postural tremor, the patients remained in a rest position first then elevated their hand, which triggered tremor almost immediately, and the tremor was present most of the time the posture was maintained. Thus postural change and tremor were present at the same-time and so the classifier may have been detecting postural activity rather than tremor *per se*. Against this, we showed that tremor frequency activities contributed most to the decoding, and in any case the distinction may be of limited clinical importance, as regardless, stimulation would be triggered during ongoing postural tremor. Second, closed-loop approaches that are based on brain signals assume that these signals do not change significantly over the long life-time of implanted electrodes. So far this has proven to be the case with regard to subthalamic LFPs in patients with Parkinson’s disease [30], but this remains to be shown in those with ET. Third, ours is essentially a technical proof-of-principal study, and the potential advantages of closed-loop DBS based on ViM thalamic LFPs remain to be established in acute and chronic clinical trials.

In conclusion, this study demonstrates that LFPs recorded from the ViM thalamus can be used to detect movements and presence of postural tremors in ET. This work lays the foundation for future work developing a closed-loop DBS system which continuously updates the decision on whether to stimulate based on activities recorded directly from the point of stimulation, in order to save battery power and minimise side effects in patients with ET.

## Materials and methods

### Participants

Seven ET patients (26-74 years old, 4 females) were recorded after obtaining informed written consent to take part in the study, which was approved by the local ethics committee. These participants underwent surgery for the implantation of DBS electrodes targeting the ViM thalamus at the Department of Functional Neurosurgery at the John Radcliffe hospital, Oxford. Leads were temporarily externalised following electrode implantation. Recordings were performed 3 to 5 days later, before final implantation and connection to the implantable pulse generator. Three patients received unilateral implantation whereas the other four patients received bilateral implantation, affording recordings from 11 ViM thalami in total. Details of the patients and the tested motor tasks are reported in Table 1.

Tremor may be considerably improved over the days following electrode implantation due to the stun effect of surgery [31,32], and prominent postural tremor was only observed in one of the seven patients recorded post-operatively in Oxford. To test whether we can decode the presence of postural tremor from thalamic LFPs, intra-operative data from another cohort of ET patients who underwent bilateral implantation of DBS electrodes into the thalamus at the Department of Stereotactic and Functional Neurosurgery in Cologne were also analysed. This afforded an opportunity to record from the micro-macroelectrodes used during target identification. The system involved electrodes of narrower diameter than the definitive DBS electrode, and, in our experience the former are less prone to induce a stun effect. Thus we could record tremor intra-operatively before implantation with the definitive DBS electrode. Raw data were first visually inspected, and those with severe artefact due to amplifier saturation or poor connection were excluded. Data sets from 12 ViM thalamus recorded from 9 patients (4 males, 67.4 ± 2.4 years old) were included for analysis in this study. The study was approved by the local ethics committee in Cologne and carried out in accordance with the Declaration of Helsinki. Detailed information about the patients and different aspects of the data have been previously described [23,33].

### Surgery procedure

The intended target coordinates were only minimally different between the two centres. For patients in Oxford, intended coordinates for the ViM thalamus were: (1) 3-5 mm posterior to the midcommissural point in the y-axis, (2) at the level of the line between the anterior and the posterior commissure in the z-axis, and (3)12-14 mm lateral from the midline (x-axis). Individual adjustments were made according to pre-operative stereotactic T2-weighted MRI so that the electrodes were 2-4 mms inside the thalamus relative to the boarder. The electrodes were placed about 3mm beyond target so that the middle contacts straddled VIM. Patients were awake during the surgery. For patients in Cologne, the ventral lateral posterior (VLp) nucleus of the thalamus was targeted on the basis of Schaltenbrand-Wahren atlas coordinates. Standard coordinates for targeting the lower border of the VLp were as follows: (1) 3-5.5 mm posterior to the midcommissural point in the y-axis, (2) at the level of the line between the anterior and the posterior commissure in the z-axis, and (3) 11.5–15.5 mm lateral to midline (x-axis).

### Experimental design

Patients in Oxford were recorded post-operatively. They were seated in a chair in front of a desktop monitor and performed different upper limb movements. In order to test the versatility of the proposed methods in detecting different movements, several motor tasks with different durations and different muscle effectors were used across different patients (see Table 1). In total, brief cued movements, such as hand gripping, joystick movement, button pressing were recorded from 5 patients (8 hands), and self-paced continuous movements, such as finger tapping, were recorded from 3 patients (5 hands). In ‘cued gripping force’ task, patients were asked to grip a dynamometer (hand dynamometer, Biometrics) so that a bar position indicating the measured force matched a cued position displayed on the monitor (Fig. 1A). Each grip lasted 6 seconds with an inter-trial interval of 4-5 seconds (randomised) and there were 25-30 trials in each session. In the finger joystick task, patient was prompted to move a joystick so that the cursor (a red dot displayed on a monitor) corresponding to the joystick position would match a target position (a green dot). For each trial, the green target jumped from the centre to one of eight potential positions, and stayed at the target position for 1 second before returning to the centre of the screen (Fig. 1B). In the ‘cued button pressing’ task, patients were asked to press the same key on a keyboard using the index finger once they saw a cue presented on a monitor. In addition, two other patients performed self-paced continuous movements. One of them (Ox4) performed blocks of continuous finger tapping; and another (Ox5) performed blocks of continuous wrist movements (extension and flexion of the wrist). Each block of movement lasted for 20-30 seconds with 20-30 seconds between the movement blocks. In order to further test the within-subject generalisability of the classifier for detecting movements, five of the seven patients performed some other self-paced movements such as spiral drawing, reaching and grasping (Table 1). Importantly, these movements were different from those used to train the classifier, so as to see if the classifier trained on pre-defined movements can decode other self-paced movements the patient might perform in everyday life.

Patients in Cologne were asked to perform a simple motor paradigm inside the operation theatre. This consisted of two conditions: (1) supine patients rested their arm in a comfortable position for 30-60 s; (2) supine patients were asked to elevate and hold their forearm contralateral to the implantation side at an angle of ~30° and to spread their fingers for 30-60 s. Subjects performed the tasks sequentially while awake after at least 15 minutes of withdrawal of sedation (remifentanil and/or propofol). Patients performed the task without speaking or performing any other activities.

### Recording

For post-operative recordings in the Oxford cohort, ViM thalamic local field potentials were recorded using a TMSi Porti amplifier (monopolar, common average reference, anti-aliasing low-pass filtering with a cut-off frequency of 500 Hz and sampling frequency of 2048Hz, TMS International, Netherlands) in patient Ox1, 2, 6 and 7. In patient Ox3-5, bipolar derivations from adjacent contact pairs were recorded through an Analog-to-digital-converter (1401power mk-II, Cambridge Electronic Design, Cambridge, UK) after amplification (Digitimer D360, Digitimer, Welfortshire, UK). Electromyography (EMG) was simultaneously recorded from the flexor and extensor carpi radialis in gripping movements, and from the first dorsal interosseous muscle (FDI) in finger joystick or finger tapping movements. Direct behavioural measurements such as generated gripping force (Biometrics hand dynamometer) or joystick positions were also simultaneously recorded using the same amplifier. In addition, 3D accelerometers (± 3g, TMSi, Netherlands) were attached to the hand in order to measure kinematic movements of the hand and the presence of tremor.

For intraoperative recordings in Cologne, a commercially available recording system (Inomed Micro Electrode Recording System; software: MER 2.4 beta) was used. Two to five micro-macroelectrodes were used, selected from a central electrode and four concentrically configured (anterior, medial, posterior, and lateral) further electrodes with a distance of 2mm from the central electrode. Local field potentials from the macroelectrodes were recorded while the electrodes were in the surgical target. Activity of the extensor digitorum communis (EDC) and flexor digitorum longus (FDL) muscles of the contralateral forearm were also simultaneously recorded using surface EMG electrodes. Both LFP and EMG signals were bandpass filtered between 0.5 and 1000 Hz during the recording and sampled at 2.5 KHz.

### Labelling of movement states based on behavioural measurements

In the Oxford cohort, direct behavioural measurements (force or joystick position) or EMG were used to identify the period of time with or without movements. EMG activities were high-pass filtered with cut-off frequency of 1 Hz, rectified and smoothed within a moving time window of 0.2 s. All behavioural measurements were normalised: gripping force was normalised to the maximal gripping force measured before the task started; the joystick position was normalised to maximal possible displacement when the joystick was displaced to its extreme; EMG activities were normalised to the 95th percentile value of the recording session. The mean and standard deviation of the background behavioural measurements during a 10 s time window with the patient at rest were quantified. For each recording session, time points with behavioural measurements over Mean + 4*STD relative to the resting baseline were labelled as ‘Movement’. All analyses were performed in MATLAB (v. 2016a, The MathWorks Inc., Natick, Massachusetts).

In the Cologne cohort, EMG activities were used to identify periods of time with or without postural tremor. EMG activities were high pass filtered with cut-off frequency of 1 Hz and rectified. Time frequency decomposition of the rectified EMG activities was obtained by applying continuous Morlet wavelet transforms with a linear frequency scale ranging from 1 Hz to 46 Hz and constant number (= 6) of cycles across all calculated frequencies. The mean and standard deviation of the peak power in the 4-7 Hz frequency band in the EMG activity when each patient was at rest were quantified. Time points in the forearm elevated blocks with EMG 4-7 Hz activity over Mean + 4*STD were labelled as ‘Postural tremor’.

### Logistic Regression based Binary Classifier

Here we adopted the logistic regression (LR) based binary (two-class) classifier. The logistic regression model predicts the probability of the presence of movements or tremor at the current time point *t* (*p(t)*) based on the linear combination of a set of predictor variables:

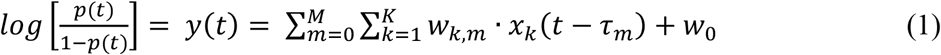

Where *K* and *M* are the number of features and the number of time lags during which the features were taken into account for the prediction, respectively, *x*_*k*_(*t* − *τ*_*m*_) is the *k*^th^ predictor variable with time lag τ_*m*_ relative to the current time point and *w*_*k,m*_ is the associated weight, *w*_0_ is an intercept constant representing the baseline probability of the occurrence of movements, and y(t) is the weighted sum of all different features. A monotonic, S-shaped continuous function (the Logistic function) was then used to map *y(t)* which ranges from −∞ to +∞ into a value between 0 and 1 (*p*(*t*)):

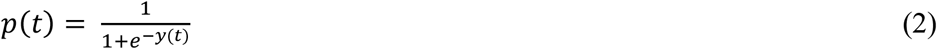

A threshold was then applied to *p(t)* to classify the current observation as movement/tremor or not.

### LFP pre-processing and feature extraction

The monopolar LFP data were re-referenced offline to obtain more spatially focal bipolar signals by subtracting the data from neighbouring electrode contacts [34]. The data were band-pass filtered between 1 Hz and 200 Hz (Butterworth filter, filter order = 4, passed forwards and backwards) and down-sampled to 1000 Hz. Time-frequency decomposition was obtained on each down-sampled bipolar channel by applying continuous Morlet wavelet transforms with a linear frequency scale ranging from 1 Hz to 195 Hz and constant number (= 6) of cycles across all calculated frequencies. Relative power was then calculated for each frequency by normalizing the absolute power by its average across time for each channel: (power average power) / average power * 100. Average movement-related modulation in the power spectra was calculated for each bipolar channel by taking the average of each 2 second epoch aligned to movement onset. The bipolar channel in each electrode with the highest modulation in the 15-35 Hz within the [−1 s, 1 s] window aligned to the movement onset (max-min) was selected for further processing. This was motivated by evidence linking maximal beta band activity and re-activity to the dorsal (motor) region of the STN [35–39].

For postural tremor detection, where LFP measurements were recorded from multiple micro macroelectrodes, the decoding was tested based on each LFP measurement. The channel with best decoding accuracy (the largest AUC value) was selected to report for that side. The best channel for decoding was the ‘central’ one for 8 out of the 12 cases, the ‘posterior’ one and the ‘medial’ one for 2 cases, and the ‘lateral’ one for 1 case.

Informed by our previous work, the power of oscillatory activities in different frequency bands over a short time window can be potential predictive features for decoding movements [40–42]. Here, the average power of eight non-overlapping frequency bands were quantified after wavelet transformation applied to the selected thalamic LFPs contralateral to the moving hand: 1-3 Hz, 4-7 Hz, 8-12 Hz, 13-22 Hz, 23-34 Hz, 35-45 Hz, 56-95 Hz and 105-195 Hz. The mean power in each of these bands was calculated over a moving time window with window length of 300 ms and overlap ratio of 40%, and then normalized against the mean power of that frequency band over the recording session. Predictive features over 10 consecutive moving windows (equivalent to 1 s preceding the current time point) were included as predictor variables. This resulted in 80 predictor variables (8 frequency bands * 10 moving windows) for the LR classifier. The output of the LR classifier was updated every 100 ms.

### Classifier training, evaluation and cross-task validation

Cross-validation was used to assess how the model, which is trained based on one training dataset, will perform when applied to a new dataset. Five-fold cross validation was performed separately for each session during which the same pre-defined movements were recorded. Five iterations of training and testing was performed on data for each recording session. In each iteration, 4 folds (80% of data) were used to train the classifier, i.e. to determine the weight *w*_*k*,*m*_ attributed to each predictor variable *x*_*k*_(*t* − τ_*m*_). The weight vector (***w***) was estimated using an optimisation function (*fminunc*) in Matlab to minimise a cost function (*j*(***w***)), which was related to mis-classification compared to the labelled state (*L*_*t*_):

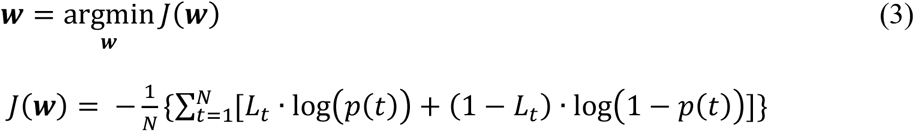

Where N is the total time points of the training data, *L*_*t*_ is the labelled movement state of each time point according to behavioural measurements, and *p*(*t*) is the output of the LR classifier of the time point given the weight vector ***w***. After the weight vector ***w*** was estimated based on the training data, the model was applied on the remaining 20% data to test the classifier performance. The training and testing were performed for five iterations so each data point was used for testing for once. The decoded movement probability reported here are concatenated ‘test’ results, with the model trained on one dataset and applied on another recorded within the same session. Subsequently, a threshold was applied to the decoded probability, to classify each time point as ‘movement’ or not. The Receiver Operating Characteristics (ROC) curve plots the true positive rate (Sensitivity) against false positive rate (1-Specificity) for different thresholds. The area under the ROC curve (AUC) provides a measure of the ability of the classifier to distinguish between the two states with 0.5 indicating a chance-level accuracy and 1 suggesting perfect classification. In this study, the ROC was plotted and the AUC was quantified to evaluate classifier performance. In addition, the detection rate and the detection latency of individual movements was also quantified. To do so, the LR classifier output around a time window between −2.5 s and +2.5 s around each individual movement onset was evaluated. A movement was treated as detected if within this time window, the LR output started from a value lower than the threshold of 0.4, increased to values higher than the threshold and stayed above this threshold for at least 500 ms. The percentage of successfully detected movements in all movements recorded in a task session was quantified as *detection rate*. The time of the LR output first exceeded the threshold relative to the actual movement onset was quantified as the *latency* of the detection.

In order to further evaluate the across-session and across-task generalisability of the LR based classifier, the classifier trained with data recorded during pre-defined movements was tested for decoding other types of self-paced movements in five patients. The ROC curves for the test were plotted; the AUC and the sensitivity (percentage of movement time that was accurately detected) were quantified and presented.

### Contribution of different LFP features in movement decoding

The percentage of contribution of LFP features in different frequency bands and different time lags (%*C*(*k*, *m*)) and the percentage of contribution from features in each frequency band (%*C*_*freq*_(*k*)) for movement or tremor decoding are calculated based on the absolute value of the weight attributed to each predictive feature (*w*_*k*,*m*_):

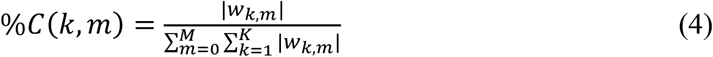

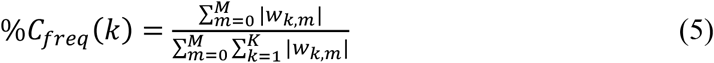

The importance of frequency bands for decoding movements was also evaluated by comparing the AUC values as the performance of the classifier after removing features of specific frequency bands.

## Acknowledgement

We acknowledge Prof. Volker Sturm, Prof. Mohammad Maarouf, Dr. Jochen Wirths and Dr. Matthias Runge for DBS-electrode placement surgery on some of the patients.

---

**Figure 3 – Figure Supplement 1.**
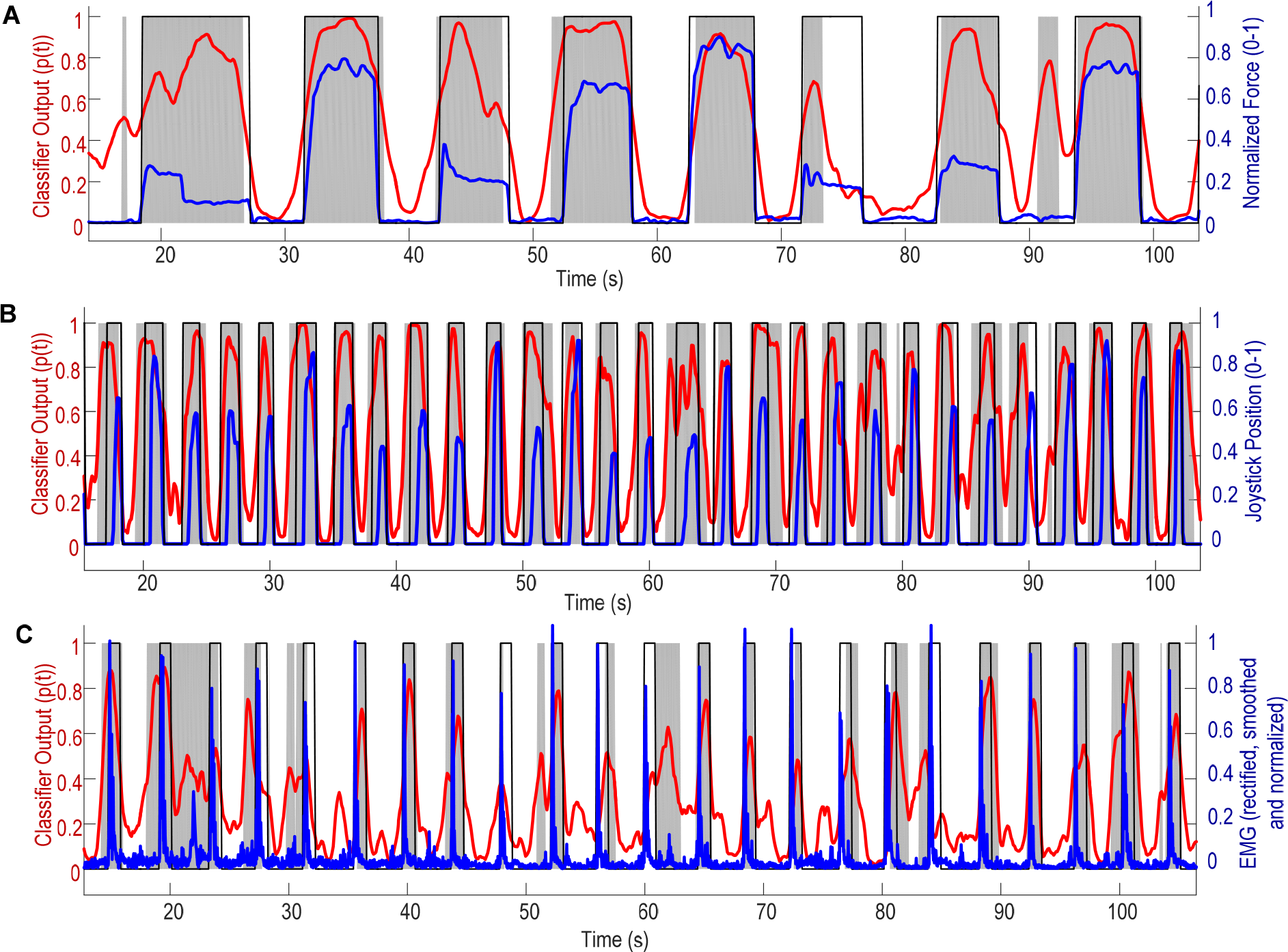
Individual brief cued movements can be decoded based on ViM LFPs. A) Hand gripping from patient Ox1; B) Finger joystick movements from patient Ox2; C) Button pressing from patient Ox3. Blue lines are behavioural measurements. These are normalized gripping force in plot A, joystick displacement relative to the centre position in Plot B, and rectified and smoothed electromyography (EMG) in Plot C. Thin black lines show the time points with movements labelled according to behavioural measurements. Red lines are the classifier output (concatenated test results from the five iterations) indicating the decoded probability of movement. The grey shaded areas are the time points when the decoded probability was above 0.4. Normalisation was to the maximum value for force and displacement, and to the 95th percentile for EMG.

**Figure 3 – Figure Supplement 2.**
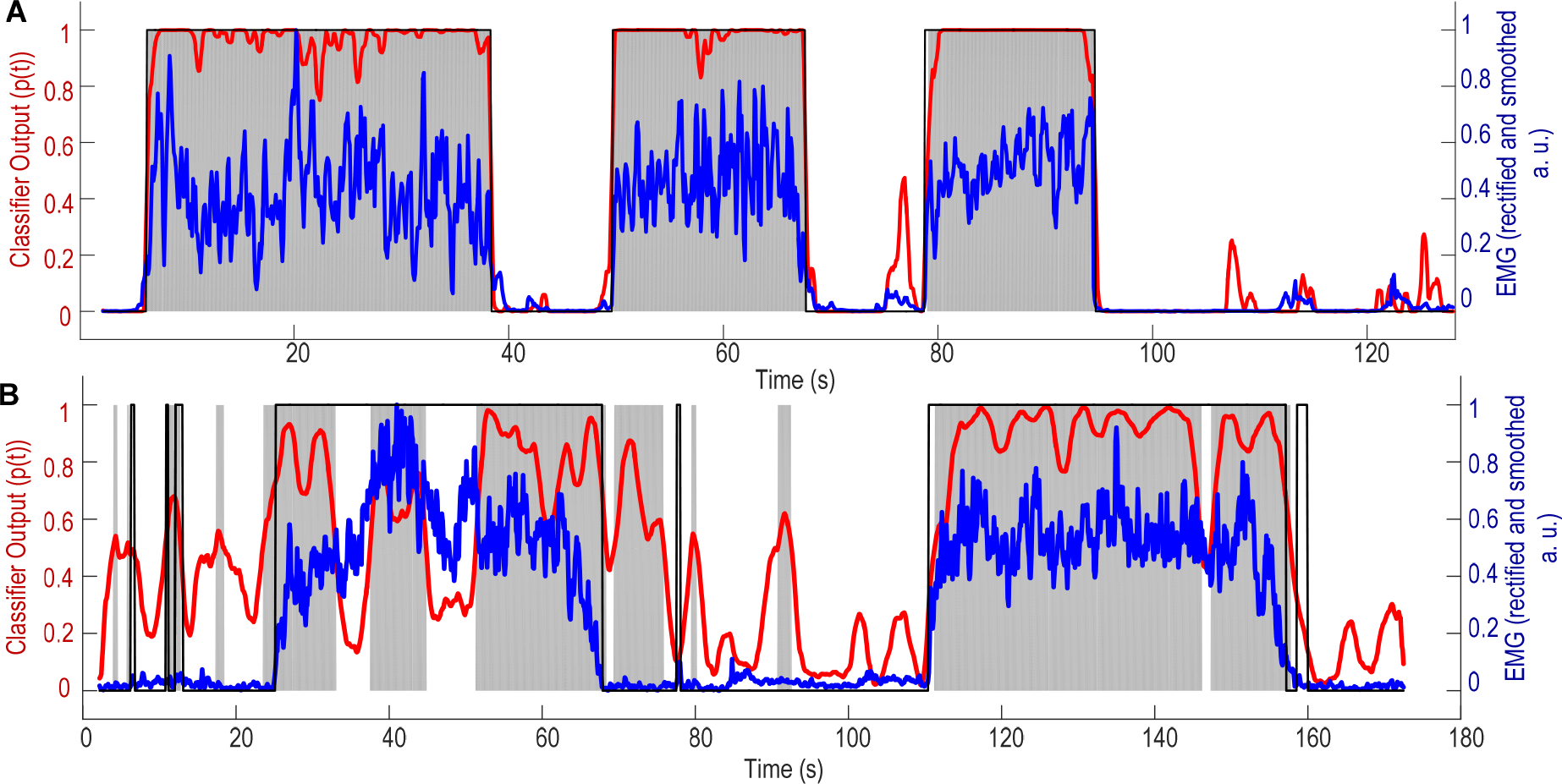
Blocks of self-paced continuous movements can be decoded based on ViM LFPs. A) Continuous finger tapping was tested in patient Ox4; B) Extension and flexion of wrist was tested inpatient Ox5. Blue lines show normalised rectified and smoothed electromyography (EMG) activity. Thin black lines show the time points with movements labelled according to EMG. Red lines are the classifier output (concatenated test results from the five iterations) indicating the decoded probability of movement. The grey shaded areas are the time points when the decoded probability was above 0.4.

